# GEMCAT – A new algorithm for gene expression-based prediction of metabolic alterations

**DOI:** 10.1101/2024.01.15.575710

**Authors:** Suraj Sharma, Roland Sauter, Madlen Hotze, Aaron Marcellus Paul Prowatke, Marc Niere, Tobias Kipura, Anna-Sophia Egger, Kathrin Thedieck, Marcel Kwiatkowski, Mathias Ziegler, Ines Heiland

## Abstract

The conclusive interpretation of multi-omics datasets obtained from high throughput approaches is an important prerequisite to understand disease-related physiological changes and to predict biomarkers in body fluids. We here present a Gene Expression-based Metabolite Centrality Analysis Tool, GEMCAT, a new genome scale metabolic modelling algorithm. GEMCAT enables integration of transcriptomics or proteomics data to predict changes in metabolite concentrations which can be verified by targeted metabolomics. In addition, GEMCAT allows to trace measured and predicted metabolic changes back to the underlying alterations in gene expression or proteomics and thus enables functional interpretation and integration of multi-omics data. We demonstrate the predictive capacity of GEMCAT on two datasets, one using RNA sequencing data and metabolomics from an engineered human cell line with a functional deletion of the mitochondrial NAD-transporter and another using proteomics and metabolomics measurements from patients with inflammatory bowel disease.

## INTRODUCTION

The amount of transcriptome data available has been increasing exponentially over the past decade. Statistical analyses and functional classification approaches like gene ontology (GO) term analyses [1, 2] are useful to generate hypotheses, but their functional interpretation is often challenging. Thus, new approaches are required that allow a better integration of expression data to predict physiologically relevant and measurable changes such as metabolic alterations. As transcriptomics, proteomics, and metabolomics become more accessible, another challenge is to effectively combine these multi-omics datasets to gain new insights into biological pathways and their regulation.

The most promising strategies for multi-omics data integration to date are based on the molecular mechanism connecting the different layers. These approaches are rooted in the understanding of gene functions and network topology [3-5]. Among these approaches, genome scale metabolic modelling is one of the most common approaches, with flux balance analysis (FBA) being the preeminent formalism. FBA is a constraint-based optimization approach and has been used extensively for biotechnological applications [5-9]. There are different approaches available to integrate gene expression data to predict metabolic flux alterations. However, FBA always requires the definition of an optimization target, like growth or ATP production, which is not always clear and easy to define, especially in mammalian systems. Although, attempts have been made to interpret results from FBA to infer metabolite concentration changes [10], the primary output of conventional FBA approaches are predictions of metabolic fluxes [9, 11, 12]. Experimental verification of metabolic fluxes in complex biological systems such as mammalian cells and whole organisms is difficult and cost-intensive, whereas direct prediction of metabolite concentration changes can be verified using targeted metabolomics and provide the basis for the prediction of metabolic biomarkers. This has been done successfully using ordinary differential equations resembling the kinetics of individual biochemical reactions [13-16]. However, due to the scarcity of kinetic data and the time required to construct these models, they are limited to a small, targeted set of reactions (up to 100) that are relevant for a specific research question [17-20]. In 2005, Patil et al. presented a network-based approach that enables the prediction of the likelihood of changes in metabolite concentrations at genome scale [21] without the necessity to define optimization targets and pathway constraints. The approach, however, cannot predict the direction of change, which would be important for the interpretation in terms of physiological consequences.

In this study, we present a novel metabolic modelling approach that enables genome-scale integration and analysis of both transcriptomics and proteomics data. In our algorithm, the genome-scale metabolic network is represented as directed graph, with metabolites as nodes and reactions as edges. Prediction of changes in metabolite concentrations is derived through the comparison of enzyme abundances between two conditions. For instance, the metabolite concentration will remain unchanged if the abundance of enzymes producing or consuming a metabolite are unchanged or all change with the same magnitude. Whereas, if the measured abundance of enzymes that produce a metabolite increase while the abundance of the ones consuming it decrease or remain constant, the concentration of this metabolite is predicted to increase. Importantly, we do not only take into account local changes in reactions directly connected to the metabolite, but also consider changes both upstream and downstream in the network. The weights of the edges are scaled based on the changes in the inferred (RNA sequencing) or measured (proteomics) enzyme abundances. Through the development of an algorithm that combines gene expression or proteomics data integration with the calculation of the PageRank (PR) centrality [22] of nodes, we can predict changes in the centrality (ranking) of the nodes within a given network. We consider the changes in the ranking equal to qualitative alterations in metabolite concentrations. An overview of our approach is shown in Figure 1. We named the approach GEMCAT – **G**ene **E**xpression based **M**etabolite **C**entrality **A**nalysis **T**ool and demonstrate its efficacy using two test cases: 1) The analysis of a transcriptome dataset from an engineered human cell line with a functional deletion of the mitochondrial NAD transporter *SLC25A51* [23], and 2) The integration of a proteomics dataset from patients with inflammatory bowel disease (IBD) [24-26]. The predictions were compared to the corresponding metabolomics data.

**Figure 1:**
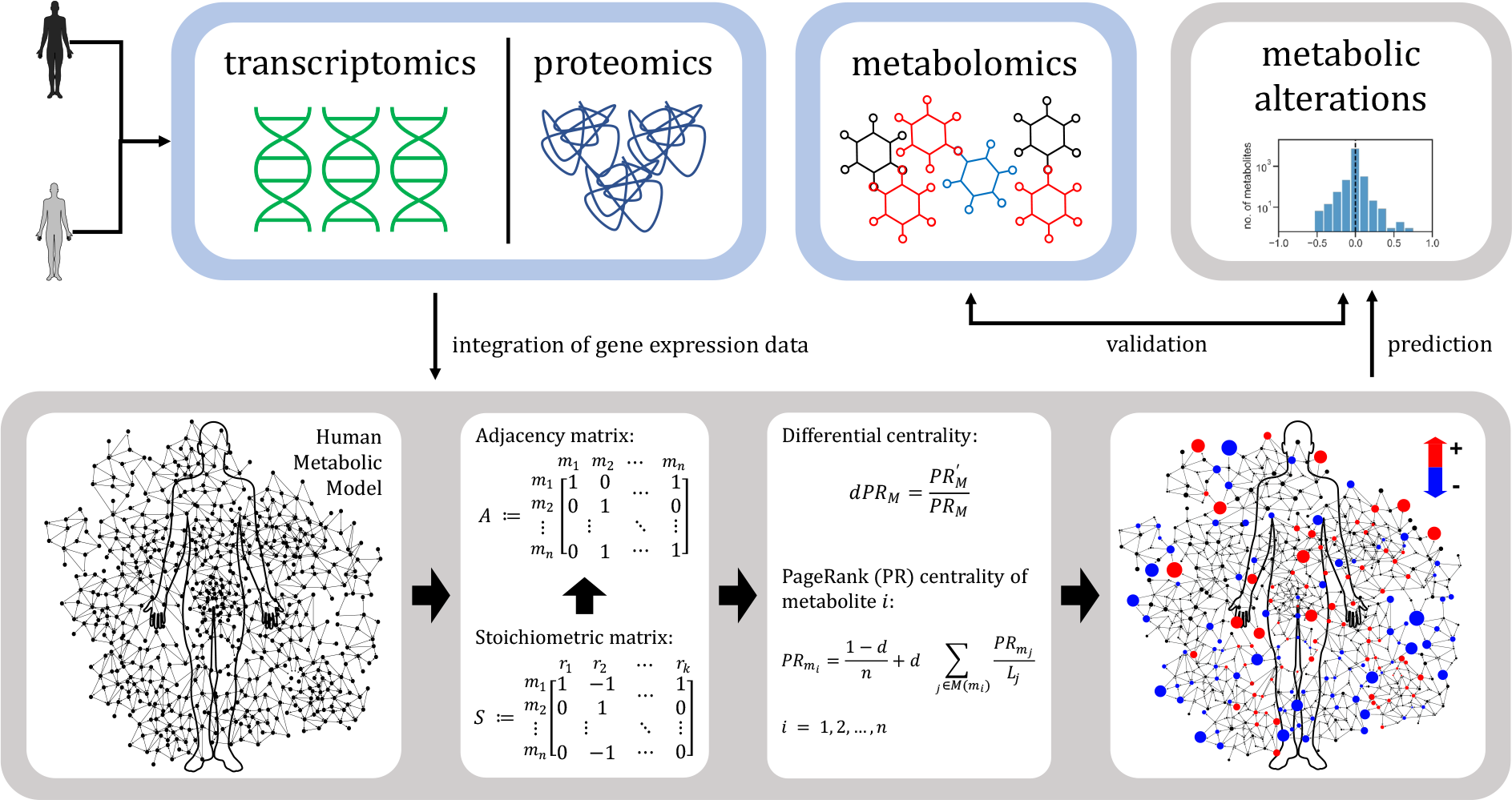
An overview of GEMCAT, our PageRank (PR) assisted method to integrate transcriptomics or proteomics data to predict metabolic alterations in human genome scale metabolic models (HMM). A stoichiometric matrix *S* and an adjacency matrix *A* can be derived from the reactions in a metabolic network. Thus, the metabolic network is represented as a directed graph composed of nodes (metabolites) linked by edges (enzymatic reactions). Upon integration of the gene expression data into the graph, PR centrality of every metabolite in the HMM is calculated. The differential PR centrality of metabolites is used to predict metabolic alteration. The predicted metabolic alterations can be validated using the experimentally measured changes in the metabolomics data.

Furthermore, to identify the expression changes that lead to certain metabolic alterations, we developed a method to calculate centrality control coefficients that links expression changes to metabolic changes and enables sensitivity analysis in genome-scale metabolic networks. This method allows us to integrate different types of omics data and perform a sensitivity analysis that can reveal the causes of metabolic alterations.

## MATERIAL AND METHODS

### Genome-scale metabolic model

We used Recon3D as a genome-scale human metabolic reconstruction that represents a comprehensive human metabolic network model, accounting for 3288 open reading frames that encode 3695 enzymes, and 13543 reactions on 8399 metabolites localised across seven subcellular locations [27].

### A graph theoretical representation of a genome-scale metabolic network

A graph is composed of nodes linked by edges. A graph is called a directed graph if the direction of the edges linking the nodes are defined. A genome-scale metabolic network can be represented as a directed graph *G*(*M,R,Φ*), where *M* is a finite set of metabolites and *R* is a finite set of interactions, which are ordered pairs of distinct interactions (*R* ⊆ {(*x,y*) | (*x,y*) *∈ M*^*2*^: *x* ≠ *y*}) contained in the metabolic network. A metabolic reaction is then represented by several edges, which is less intuitive, but makes the graph easier to process as the nodes represent the same kind of entities [28]. This approach allows the integration of information about the metabolites transition on edges. Several ways of representing a metabolic network as a graph exist [28, 29]. A stoichiometric matrix *S* := (*s*_*ij*_), where rows represent metabolites and columns denote reactions can be used to derive a metabolic graph. *s*_*i,j*_ takes negative integer for metabolites that are substrates and positive integers for products of a reaction [19]. The elements are zero otherwise. *S* can be transformed to yield an adjacency matrix *A* = (*a*_*ij*_), where *a*_*i,j*_ gets 1 when a reaction exists between metabolite *m*_*i*_ ∈ *M* and *m*_*$*_ ∈ *M* (*i* ≠ *j*). The entries are zero otherwise. The edges are characterised by the weighted adjacency matrix *Φ* = (*ϕ*_*i,j*_), where *ϕ*_*i,j*_ is the sum of weights of the edges between metabolite *m*_*i*_ and *m*_*j*_. We derived *ΔΦ* from the differential abundance of enzymes by comparing two different strains or conditions.

### Integration of the differential gene expression or proteomics data

The integration of differential measured (proteomics) or inferred (transcriptomics) protein abundance starts with the mapping of the corresponding genes or proteins onto the metabolic network. This is done by processing the gene-protein-reaction (GPR) relations described by Fang et al. [30]. According to which, the following scenarios are possible: a) if only one protein is associated with a metabolic reaction, the abundance of this protein is assigned to the reaction; b) if several proteins are jointly required for a reaction to take place, the geometric mean of the abundance is assigned the reaction; c) if any one of several proteins is sufficient for a reaction to occur, the arithmetic mean of the abundance is assigned to the reaction; and d) any GPR containing combinations of both b and c is parsed according to their logical relation. The processing of GPRs results in an abundance vector, where each element denotes the net abundance of a protein catalysing a particular reaction.

### Calculation of the weighted adjacency from a stoichiometric matrix

Let *E* be the protein abundance vector calculated by processing the corresponding GPRs. *S* be the stoichiometric matrix of the given metabolic network with rows and columns representing metabolites and reactions, respectively. All reversible reactions contained in *S* were split into their unidirectional component reactions, and the corresponding entry in *E* was duplicated accordingly. A weighted stoichiometric matrix is created as

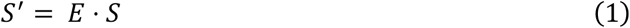

We split this matrix into its components for products and reactants, respectively, as

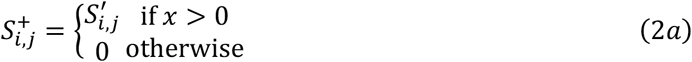

And

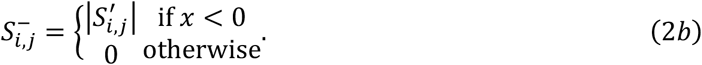

These matrices are used to calculate the weighted adjacency matrix as

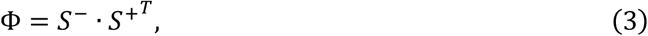

which is a square matrix used to represent a finite graph, which in our case is the human metabolic network from Recon3D model. The elements of the matrix indicate the weight shared by the adjacent pairs of metabolites in the metabolic network.

### Calculation of the differential centrality to predict metabolic alterations

The PageRank (PR) centrality of metabolites *M* can be calculated as

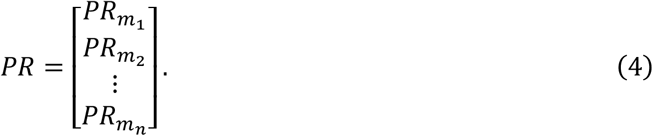

The PR centrality provides a ranking which identifies the importance of a node in a network. In its original form, the algorithm calculates a probability distribution to represent the likelihood of a person randomly clicking on links and arriving at a particular page on the internet [22, 31]. This probability is expressed as a numeric value between 0 and 1. In the context of a metabolic network, the PR value for any metabolite *m*_*i*_ is dependent on the PR values for each metabolite *m*_*j*_ (*i* ≠ *j*) contained in the set *M*(*m*_*i*_) containing all metabolites linking to metabolite *m*_i_ divided by the total number of links from metabolite *m*_*$*_. Further, metabolites with no outbound edges i.e., metabolites that are sinks, are assumed to link out to all other metabolites in the network. Their PR scores are therefore divided evenly among all other metabolites. There is a residual probability *d*, also known as damping factor, that is added to each node as a likelihood of a transition to any other node in the network [22]. The PR value of a metabolite *m*_*i*_ can be calculated as

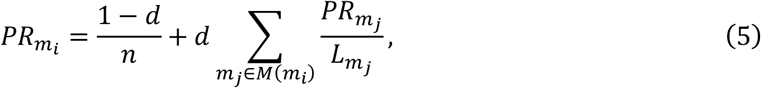

where 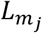 is the number of outbound links on *m*_*j*_, and *n* is the total number of metabolites in the metabolic network. The PR values correspond to the elements of the dominant right eigenvector of the weighted adjacency matrix, which has been normalised to ensure that each column adds up to one. The eigenvector is given in Equation 4. The PR centralities are iteratively calculated for each metabolite until the convergence is achieved [32]. A differential PR centrality (*dPR*) can be calculated by comparing the relative change in the PR centralities of *M* for the perturbed and baseline conditions. We used *dPR* to predict changes in the abundance of *M* (Equation 6).

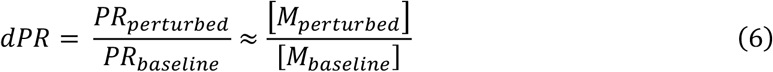

### Gene-expression and metabolomics measurements

Cells (293-*SLC25A51*-ko and parental HEK293) were cultivated in DMEM-high glucose (Merck/Sigma D5671) supplemented with 10% (v/v) FBS, 2 mM L-glutamine, 1 mM sodium pyruvate and penicillin/streptomycin at 37°C in humidified atmosphere with 5% CO_2_. Cells were harvested, washed with PBS and counted. For RNA-Seq analyses 5 × 10^6^ cells in 3 technical replicates per cell line were frozen in liquid nitrogen and shipped on dry-ice to Novogene Co., Ltd, Cambridge, UK for processing. RNA sequences were generated using the Illumina NovaSeq platforms. Fragments per kilobase of transcript per million mapped reads (FPKM) were used directly to infer enzyme abundance changes. For the metabolomics measurements, five times 2 million cells from each parental HEK293 and 293-*SLC25A51*-ko cells were subjected to chloroform-methanol based simultaneous proteo-metabolomics liquid-liquid extraction (SPM-LLE) [33]. Glycolytic metabolites, metabolites of tricarboxylic acid (TCA) cycle and nucleotides (ATP, ADP, AMP) were analysed by ion chromatography-single ion monitoring-mass spectrometry (IC-SIM-MS) [33] using a quadrupole orbitrap (Exploris 480) and an ICS-6000 IC system (both Thermo Fisher Scientific). Free amino acids and tryptophan metabolites were analysed by multiple reaction monitoring (MRM) using a triple quadrupole mass spectrometer (TQ-XS, Waters) coupled to an UPLC-system (ACQUITY Premier, Waters) as described previously [34]. IC-SIM-MS data was processed using TraceFinder 5.0 (Version 5.0.889.0, Thermo Scientific) and the LC-MRM-MS data was processed using MS Quan (Waters Connect, Waters) [33, 34]. The differential proteomics and metabolomics from mucosa biopsies of IBD patients and healthy controls were obtained from [24-26].

### Chemicals

HPLC-grade acetonitrile (ACN), methanol (MeOH), formic acid (FA), as well as Micro BCA Protein Assay Kit, Gibco Qualified FBS, and ammonium bicarbonate were obtained from Thermo Fisher Scientific (Dreieich, Germany). [U-13C]-labelled yeast extract (2 billion cells) was purchased from ISOtopic Solutions (Vienna, Austria), reconstituted in 2 mL HPLC-H20, aliquoted and stored at −80 °C. [U-13C]-labelled lactate (20 % w/w dissolved in H_2_O) and [U-13C-15N]-labelled canonical amino acids (dissolved in 0.1 M HCl) were purchased from Eurisotop (Saarbruecken, Germany).

## RESULTS

### Metabolic alterations in an engineered HEK293 cell-line

We used RNA-Seq data from an *SLC25A51* CRISPR-Cas9 knockout cell line that lacks a functional mitochondrial NAD transporter [23]. Differential gene expression was calculated by comparing the RNA-Seq from three different replicates of *SLC25A51* knockout (293-*SLC25A51*-ko) and parental HEK293 cells respectively. We used Recon3D [27] as a genome-scale human metabolic model (HMM). 2615 transcripts could be mapped to the Recon3D model. As the model contains 3695 enzymes that catalyse 7675 out of 13543 reactions in total, we assumed that the remaining reactions did not change and thus set the corresponding values to 1. Truncated transcripts of *SLC25A51* can still be detected in the 293-*SLC25A51*-ko cell line, which is common in CRISPR-Cas9 deletions. However, the massive decrease of mitochondrial NAD has been shown experimentally, thereby functionally validating the knockout [23]. We thus set the transport reaction of NAD across the mitochondrial membrane to 0. We used our newly developed algorithm GEMCAT to calculate the differential PageRank centrality (*dPR*) of metabolites *M* (*M* = *m*_1_,*m*_*2*_,…,*m*_*n*_: *n* = 8399) in the HMM and used *dPR* as an estimate for metabolic alteration that is caused by changes in transcript abundance in 293-*SLC25A51*-ko compared to parental HEK293 cells. The predicted log2fold changes vary between -0.75 and 0.75 (see Figure 2a). As Recon3D is a compartmentalized model but we only have whole cell metabolomics measurements, we calculated the arithmetic mean of predicted metabolite changes between three subcellular compartments—cytosol, mitochondria, and nucleus (compartment specific predictions are shown in Supplementary Figure S1). We compared the mean values with experimentally measured concentration changes in 43 metabolites. 29 out of 43, thus ∼67% of the metabolites, were predicted correctly (see Figure 2b). This includes several intermediates of the TCA cycle that have previously been reported to decrease in *SLC25A51*-deficient HAP1 cells [35]. Further details about variation between replicates are provided in Supplementary Figures S1, S2 and Table S1.

**Figure 2:**
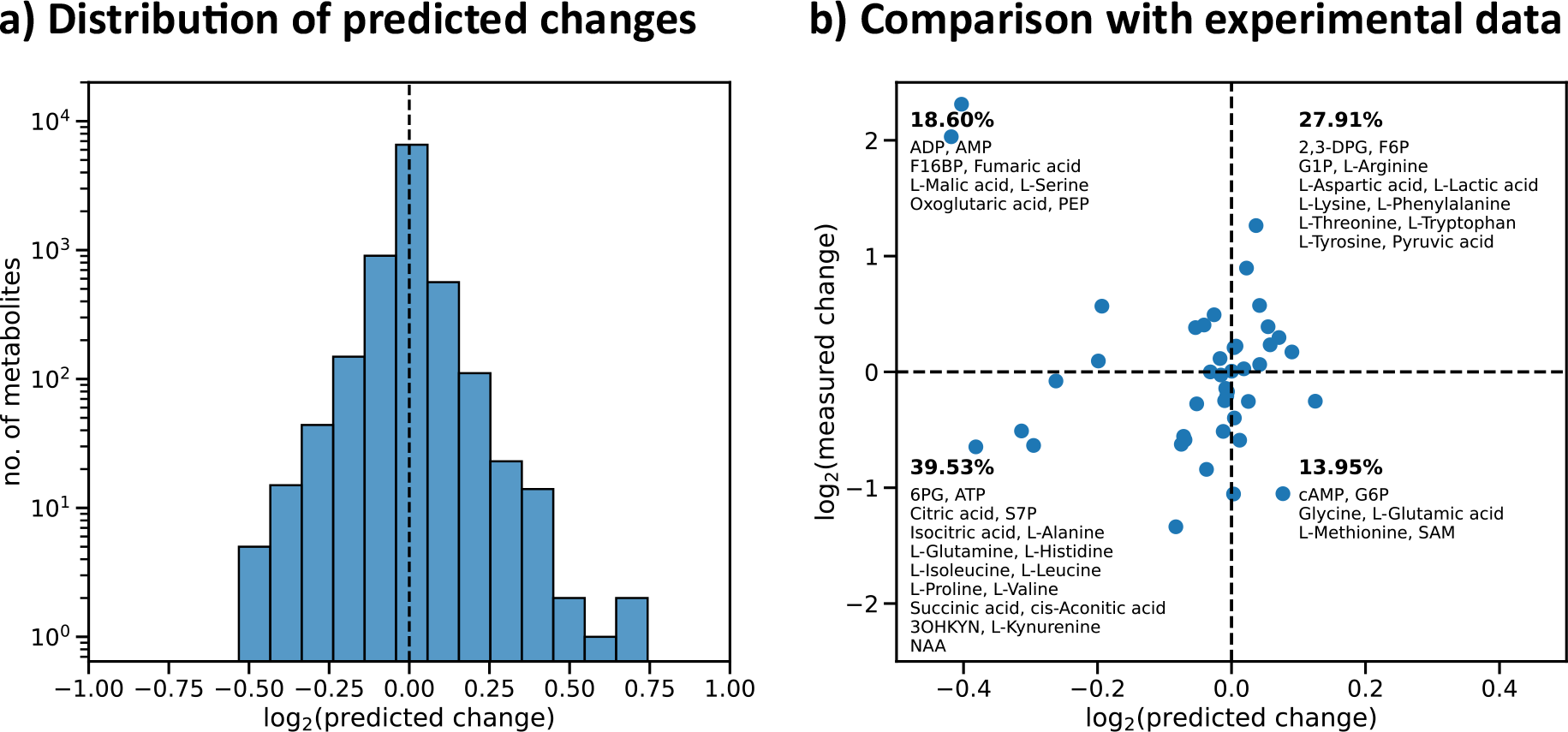
Comparison of predicted and measured changes in the abundance of metabolites in *SLC25A51*-deficient (293-*SLC25A51*-ko) cells relative to parental HEK293 cells. (a) A histogram showing the distribution of changes predicted in the abundance of 8399 metabolites in the HMM (Recon3D model) using RNA-Seq data from three replicates. (b) A scatter plot showing the distribution of the mean of predicted metabolic changes calculated based on the integration of RNA-Seq data from three replicates in comparison to the mean of the experimentally measured changes in metabolite concentrations in 293-*SLC25A51*-ko relative to parental HEK293 cells from five replicates (for details see Materials and Methods). The dashed black lines are used to divide the plot into four quadrants. The upper right and lower left quadrants show metabolites, whose concentration changes are predicted correctly. In each quadrant the percentage and names of metabolites corresponding to it are shown. Metabolite abbreviations: AMP: Adenosine monophosphate; ATP: Adenosine triphosphate; cAMP: Cyclic AMP; 2,3-DPG: 2,3-Diphosphoglyceric acid; 6PG: 6-Phosphogluconic acid; S7P: D-Sedoheptulose 7-phosphate; F16BP: Fructose 1,6-bisphosphate; F6P: Fructose 6-phosphate; G1P: Glucose 1-phosphate; G6P: Glucose 6-phosphate; PEP, Phosphoenolpyruvic acid; SAM: S-Adenosylmethionine; NAA: N-Acetyl-L-aspartic acid; 3OHKYN: Hydroxykynurenine.

### Predicting metabolic alterations in IBD patients

To predict metabolic alterations in IBD patients with ulcerative colitis (UC), we used a published mucosa proteome dataset comparing protein abundance in patients with severe UC to that of healthy adults [26]. The dataset comprises only significantly changed protein abundances.

These were mapped to 158 enzymes in the Recon3D model. The values for the enzymes not covered by the proteomics were set to 1 and thus assumed to be unchanged between patients and controls. Upon integration of the differential proteomics data, we calculated the *dPR* scores of all metabolites in our HMM to predict the changes in their concentrations. The distribution of predicted changes in the metabolite concentrations was centred around 0 (see Figure 3a) with most of the metabolites being unchanged as expected as very few enzymes in the network were significantly changed. To validate our predictions, we compared them with the metabolite measurements from the same set of UC patients and healthy adults [24, 25]. We correctly predict changes in ∼79% of the metabolites measured to change significantly (i.e., *p <* 0.05 [25], cf. orange dots in Figure 3b). See Supplementary Figure S3 for the comparison of all measured metabolites and Table S2 for values and standard deviation.

**Figure 3:**
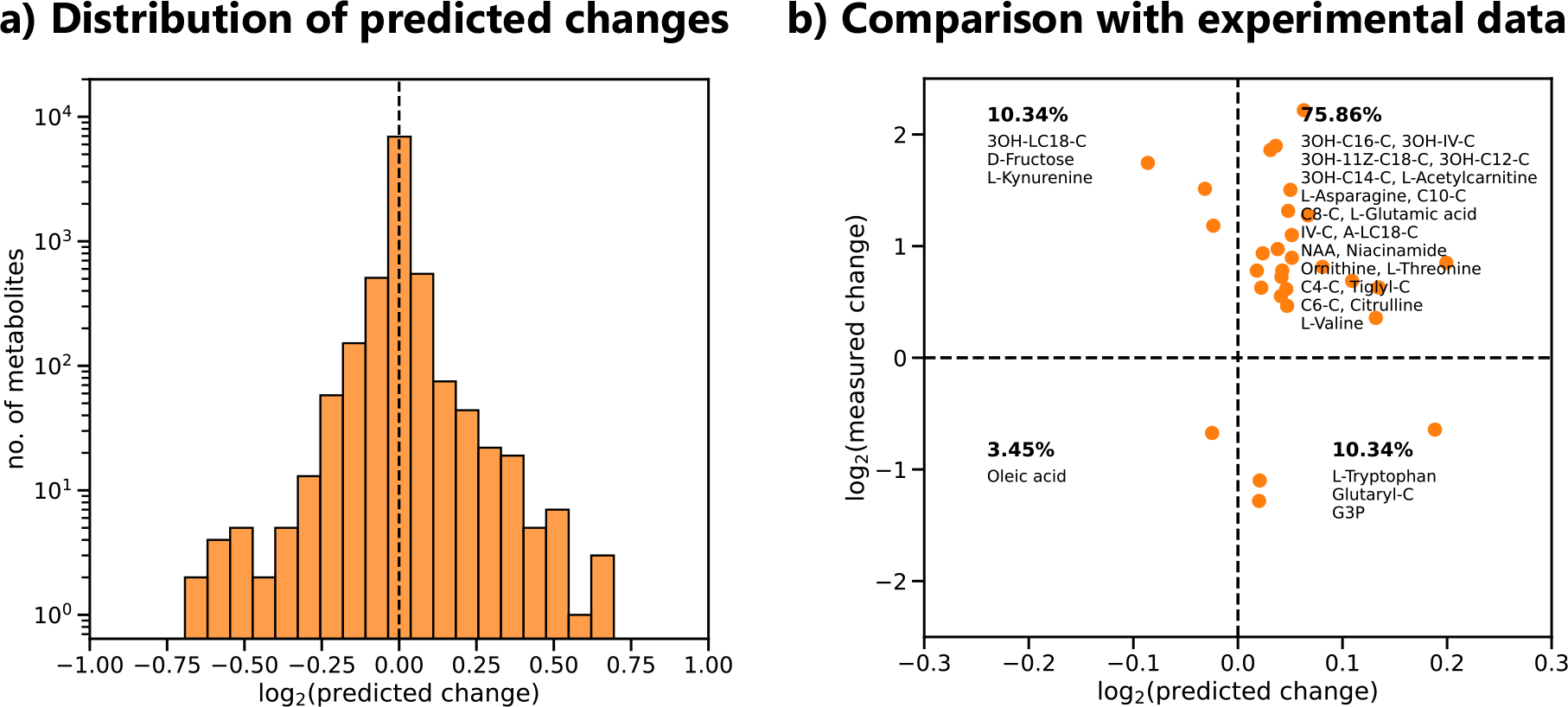
Comparison of predicted and measured changes in the abundance of metabolites in UC patients relative to healthy controls. (a) The histogram shows the distribution of predicted change in all 8399 metabolites contained in the HMM. (b) A scatter plot showing the comparison of the predicted changes against the experimentally measured metabolic changes. The dotted black lines separate the quadrants between correctly (upper right and lower left quadrants) and incorrectly predicted metabolites. All metabolites measured to be changed significantly (*p* < 0.05) and with *dPR* > ‖0.014‖ are shown. The complete results for 137 metabolites are provided in Supplementary Figure S3 and Table S2. Metabolites are shown in orange and corresponding names are indicated in the respective quadrants. The percentage of metabolites in each quadrant is also shown. Metabolite abbreviations: 3OH-C16-C: 3-Hydroxyhexadecanoylcarnitine; 3OH-IV-C: 3-Hydroxy-Isovaleryl Carnitine; 3OH-11Z-C18-C: 3-Hydroxy-11Z-octadecenoylcarnitine; 3OH-C12-C: 3-Hydroxydodecanoylcarnitine; 3OH-C14-C: 3-Hydroxy-Tetradecanoyl Carnitine; C10-C: Decanoylcarnitine; C8-C: L-Octanoylcarnitine; 3OH-LC18: (3S)-3-Hydroxylinoleoyl-CoA; IV-C: Isovaleryl Carnitine; NAA: N-Acetyl-L-aspartic acid; A-LC18-C: Alpha-linolenyl carnitine; C4-C: Butyrylcarnitine; Tiglyl-C: Tiglyl Carnitine; Glutaryl-C: Glutaryl Carnitine; C6-C: Hexanoyl Carnitine; and G3P: Glycerol 3-phosphate.

### Multi-omics integrations allow tracing of metabolic changes to changes in enzyme abundance

As the accessibility of metabolomics measurements increases, it becomes important to integrate them with other omics datasets such as transcriptomics and proteomics. We therefore extended GEMCAT to enable tracing of metabolic changes back to the changes in gene expression or proteomics. To this end, we first analysed the impact of abundance changes in each enzyme on the predicted change of all metabolites. This approach is similar to metabolic control analysis and is an important component of mathematical analysis of complex systems [36, 37]. It describes the system behaviour in terms of the properties of its variables. To systematically analyse the effect of perturbation of the metabolic network, we estimated the centrality control coefficient 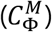 of metabolites *M* due to the perturbation in *Φ*. The control on the centrality of a metabolite *m* by an enzyme *x* was calculated as

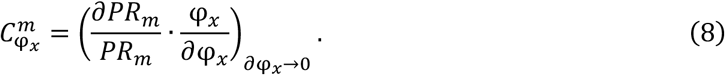

If this coefficient is positive the PR centrality of the metabolite *m* increases as the weight of the reaction increases, which in turn is increased due to the increase in abundance of enzyme *x*, and *vice-versa*. These coefficients are characteristic for a given network and are not dependent on the expression data. Furthermore, 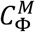 can be used to estimate the change in centrality of *M* for a given differential in enzyme abundance 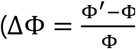, where *Φ*′ and *Φ* denote enzyme abundances for compared and reference condition, respectively) and refer to this as response coefficient 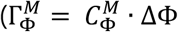; Equation 8). The response coefficients 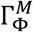 can be used to trace metabolic alterations back to the underlying changes in the gene expression or proteomics as shown in Figure 4. (The full set of centrality control coefficients 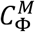 and response coefficients 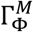 are provided as Supplementary Tables S3 and S4 uploaded to https://github.com/MolecularBioinformatics/prm_manuscript.) Alternatively, the approach can be used to analyse discrepancies between predictions and experiments and thus identify potential points of e.g. post-translational regulation or potential incomplete pathway information.

**Figure 4:**
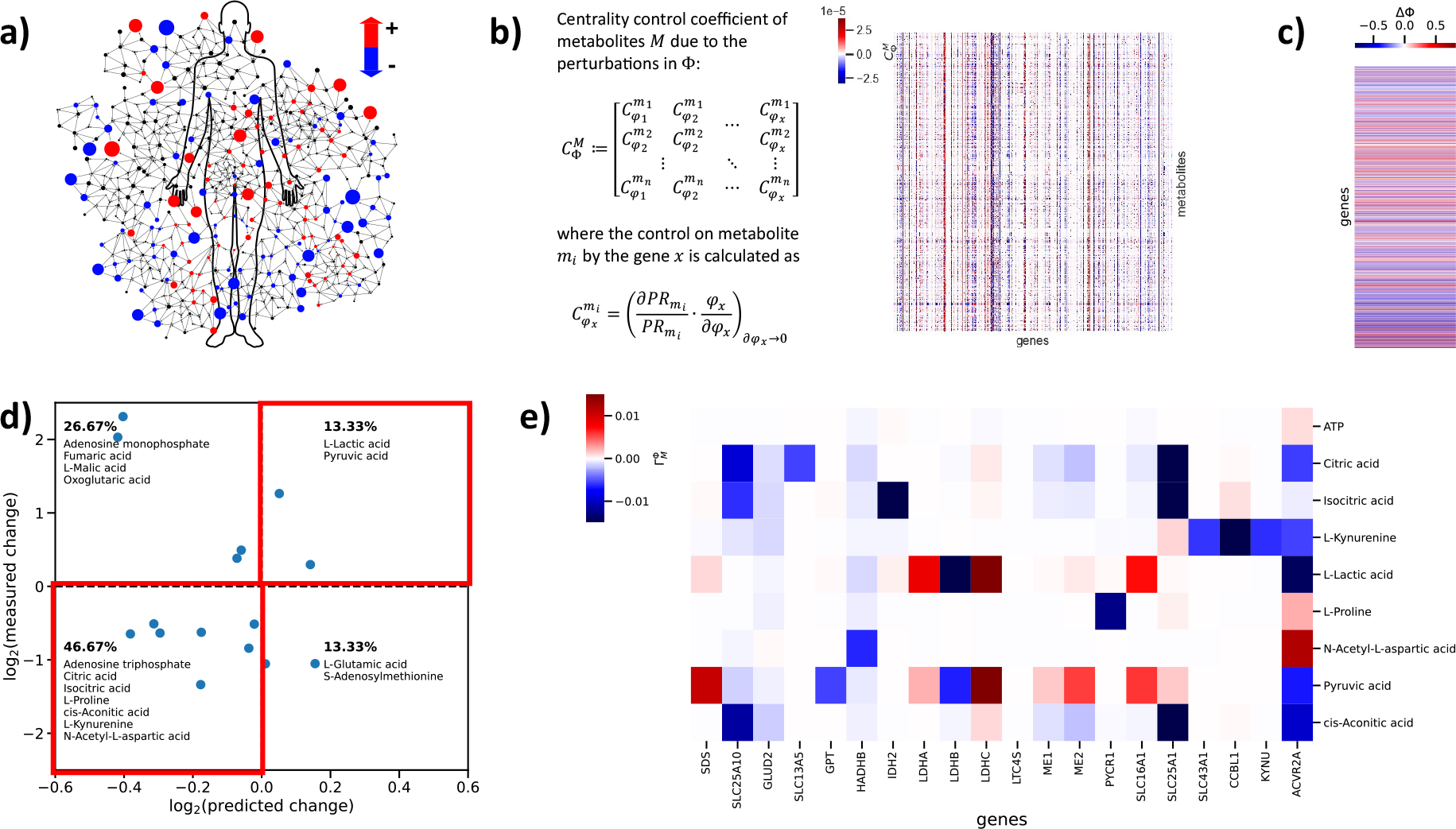
Calculation of response coefficients 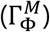 to trace metabolic alterations back to the underlying changes in the gene expression or proteomics. (a) Changes in metabolomics data mapped onto the human metabolic network. (b) Calculation of centrality control coefficients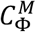. A heatmap of 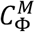 corresponding to 8399 metabolites and 2795 metabolic genes is shown. (c) A heatmap of a differential gene expression (*ΔΦ*), which is calculated by comparing the RNA-Seq data of *SLC25A51*-deficient and parental HEK293 cells). (d) The scatterplot is based on the comparison shown in Figure 2b but limited to metabolites that show consistent predicted changes for three replicates of RNA-Seq data and significant changes in the measurements (*p <* 0.05, two-tailed Student’s t-test). For details see Supplementary Figure S2b. Each dot represents the mean value of a metabolite. The red bold lines highlight the upper-right and bottom-left quadrants, where the direction of the predicted changes agree with the experimentally measured changes. In each quadrant the percentage and names of metabolites corresponding to it are shown. (e) Calculation of response coefficients 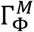 for a given differential gene expression (*ΔΦ*). A heatmap showing the response coefficients for correctly predicted metabolites corresponding to the genes with highest response coefficients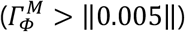.

## DISCUSSION

In this study, we present GEMCAT, a new algorithm to predict metabolic alterations in a large set of metabolites using genome-scale metabolic networks. Unlike FBA, GEMCAT does not require an objective function, and only uses the intrinsic properties of the metabolic network. Furthermore, it can predict qualitative changes in metabolite concentrations directly. Upon integration of enzyme abundances into a graph-based representation of the human metabolic network, we calculated the change in PR centrality of metabolites contained in the graph. This change in PR centrality, referred to as differential PR (*dPR*) centrality in this paper, was used to predict metabolic alterations originating from changes in enzyme abundances. Using Recon3D as human genome scale metabolic reconstruction, GEMCAT can predict changes in 8399 metabolites for seven sub-cellular compartments. As metabolomics measurements are limited to a much lower number of metabolites and are difficult, if not impossible, to perform for all subcellular compartments, we can only verify our predictions on a limited set of metabolites. Since most metabolomics data refer to whole-cell analyses, we had to combine the predicted subcellular changes into whole cell changes, and here used a simplified assumption i.e., the calculation of arithmetic means of subcellular changes. To accurately determine the contribution of each subcellular compartment to changes at the whole cell level, compartment-specific concentrations of each metabolite would be required. Such data is not available to date. Compared to transcriptomics, proteomics data are considered a better proxy for enzyme activities. But, the protein coverage is still much lower in proteomics compared to transcriptomics datasets. Despite these limitations, GEMCAT achieves a remarkably high prediction accuracy for the patient dataset even though the dataset only covered 158 enzymes of the network. Given that untargeted metabolomics measurements are still rather expensive and incomplete, GEMCAT provides a new approach to derive hypotheses about metabolic alterations that can be validated by far more accurate targeted metabolite measurements. In this way, GEMCAT can also assist in the identification of disease-specific metabolic biomarkers or signatures.

To understand the reasons for incorrectly predicted metabolic changes we traced them back to the network representation of the corresponding pathways. We recognized, for example, that Recon3D includes only a limited number of reactions that are sinks for methyl-group donors such as S-adenosyl methionine (SAM), which has an impact on the prediction accuracy for SAM as well as for serine and methionine, serving as methyl-group sources. Consequently, our approach can be applied to improve metabolic network reconstruction by tracing incorrectly predicted metabolites to identify errors or gaps in the network reconstructions. Finally, we envisage extending GEMCAT to include kinetic and other relevant information such as post-translational modifications and their functional effects to adjust the weighting of the edges and further improve the predictions.

## Supporting information

Supplementary Figure S1, S2 and S3 as well as Supplementary Tables S1 and S2

## DATA AVAILABILITY

A computational framework to allow GEMCAT calculations is developed using python programming. The framework is available as a python package and can be downloaded from Python Package Index (PyPI). GEMCAT is also hosted on GitHub (https://github.com/MolecularBioinformatics/GEMCAT) and has been provided via figshare (https://doi.org/10.6084/m9.figshare.25020101.v1). The Supplementary Tables (S3 and S4) as well as the differential expression data set, the proteomics and metabolomics data along with the python scripts used to produce images presented in this paper can be downloaded from here (https://github.com/MolecularBioinformatics/prm_manuscript). The RNA sequencing data is available at GEO GSE255209.

## SUPPLEMENTARY DATA

Supplementary Figure S1, S2 and S3 as well as Supplementary Tables S1 and S2 have been combined into one file and uploaded alongside the manuscript.

