## Supplementary Figure S1, S2 and S3 as well as Supplementary Tables S1 and S2 for "GEMCAT – A new algorithm for gene expression-based prediction of metabolic alterations"

### FIGURES

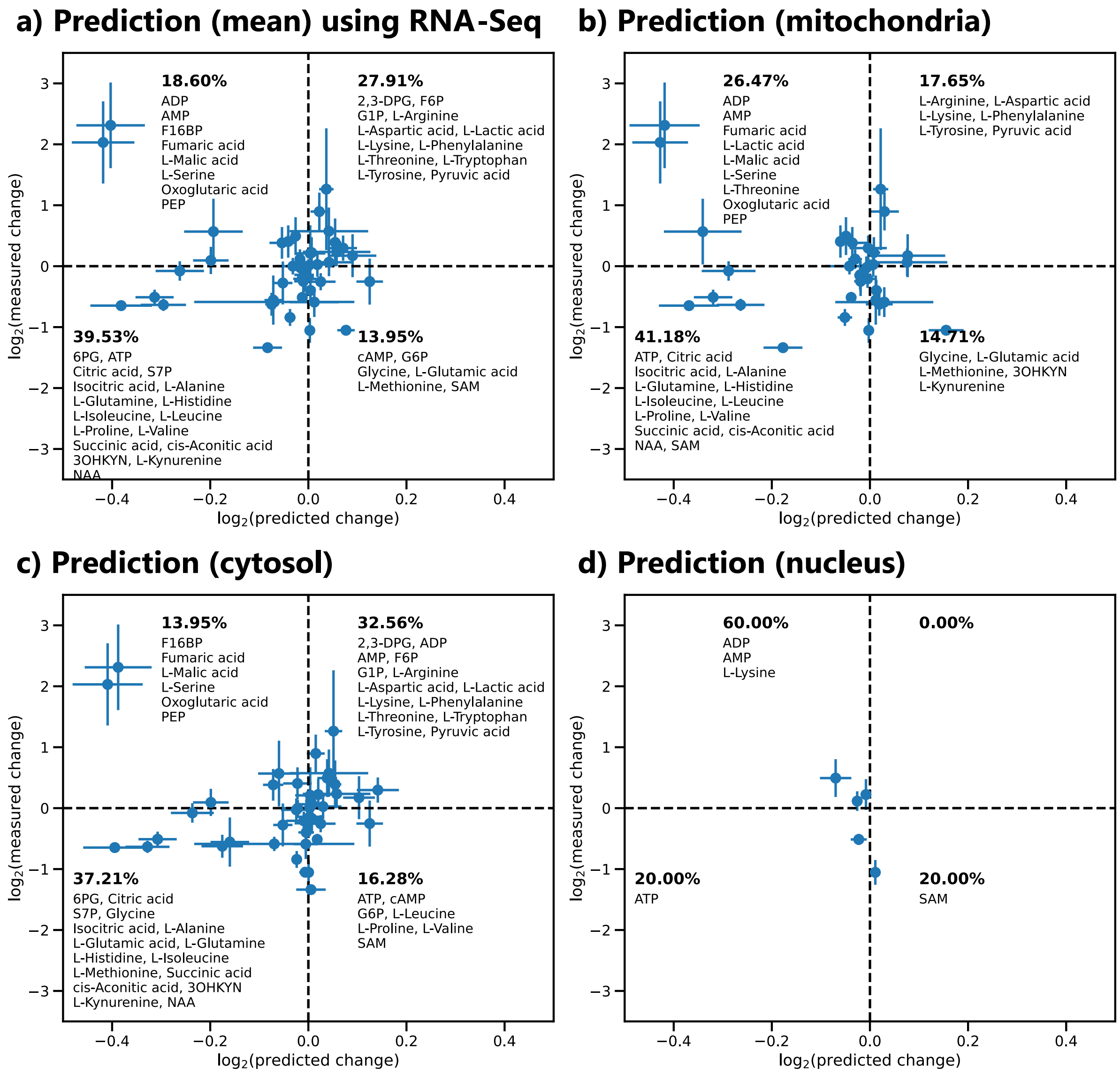


Figure S1: Comparison of predicted and measured changes in the abundance of metabolites in *SLC25A51*-deficient (293-SLC25A51-ko) cells relative to parental HEK293 cells. The dashed black lines separate the quadrants between correctly (upper right and lower left quadrants) and incorrectly predicted metabolites. Each data point represents the arithmetic mean of the predicted changes, whereas the bars represent standard deviation calculated based on individual RNA-Seq data. a) A scatter plot showing the comparison of the mean of the changes predicted across replicates and three subcellular locations (cytosol, mitochondria, and nucleus) against measured changes in the metabolites. b) A scatter plot showing the mean of predicted changes from three replicates in the metabolites localised in mitochondria only against experimentally measured changes in metabolites. c) A scatter plot showing the mean of predicted changes from three replicates in the metabolites localised in cytosol only against experimentally measured changes in metabolites. d) A scatter plot showing the mean of predicted changes from three replicates in the metabolites localised in nucleus only against experimentally measured changes in metabolites. All predicted changes are based on the integration of individual RNA-Seq from three replicates (fold changes calculated from FPKM for all transcripts), and metabolomics measurements from five replicates. Each dot represents a metabolite, and metabolite names and the percentage of analysed metabolites are indicated in the respective quadrants. Metabolite abbreviations see Figure 2.

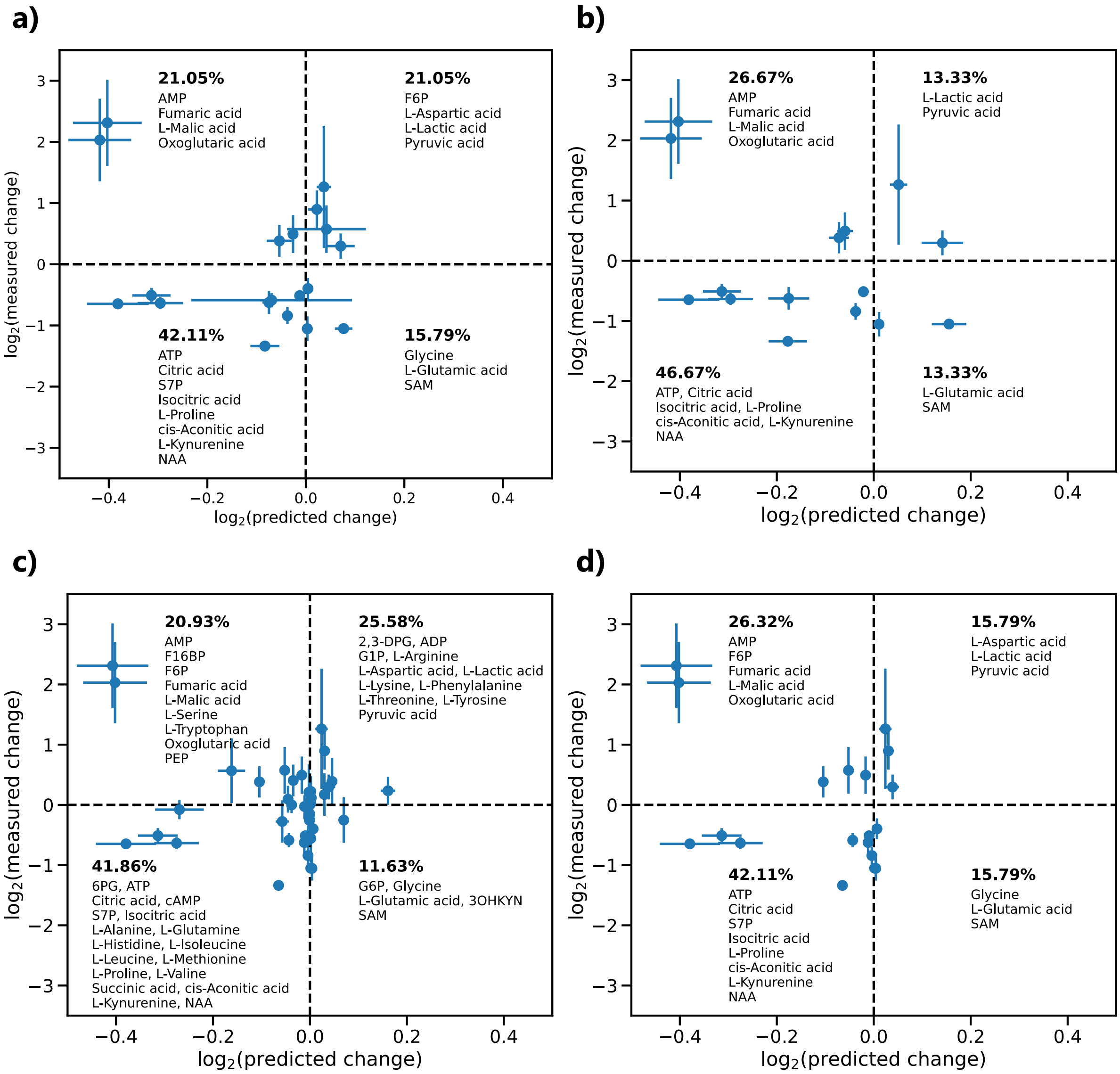


Figure S2: Comparison of predicted and measured changes in the abundance of metabolites in *SLC25A51*-deficient (293-SLC25A51-ko) cells relative to parental HEK293 cells. The dashed black lines separate the quadrants between correctly (upper right and lower left quadrants) and incorrectly predicted metabolites. Each dot represents the arithmetic mean of the predicted changes across three subcellular compartments i.e., cytosol, mitochondria, and nucleus calculated based on RNA-Seq data integration. Whereas the bars represent standard deviation calculated based on individual RNA-Seq data. a) A scatter plot showing the predicted changes calculated based on the integration of RNA-Seq from three replicates in comparison to experimentally measured significant ($p < 0.05$, two-tailed Student's t-test) changes in metabolite concentrations in 293-SLC25A51-ko relative to parental HEK293 cells from five replicates. b) A scatter plot showing the comparison of consistently predicted metabolic changes calculated based on the integration of RNA-Seq data from three replicates in comparison to the experimentally measured significant changes ($p < 0.05$, two-tailed Student's t-test) in metabolite concentrations in 293-SLC25A51-ko relative to parental HEK293 cells from five replicates. c) A scatter plot showing the predicted changes calculated based on the integration of significantly changed ($p < 0.05$, two-tailed Student's T-test) RNA-Seq from three replicates in comparison to experimentally measured changes in metabolite concentrations in 293-SLC25A51-ko relative to parental HEK293 cells from five replicates. d) A scatter plot showing the predicted changes calculated based on the integration of significantly changed ($p < 0.05$, two-tailed Student's T-test) RNA-Seq from three replicates in comparison to experimentally measured significant changes ($p < 0.05$, two-tailed Student's t-test) in metabolite concentrations in 293-SLC25A51-ko relative to parental HEK293 cells from five replicates. Metabolite abbreviations see Figure 2.

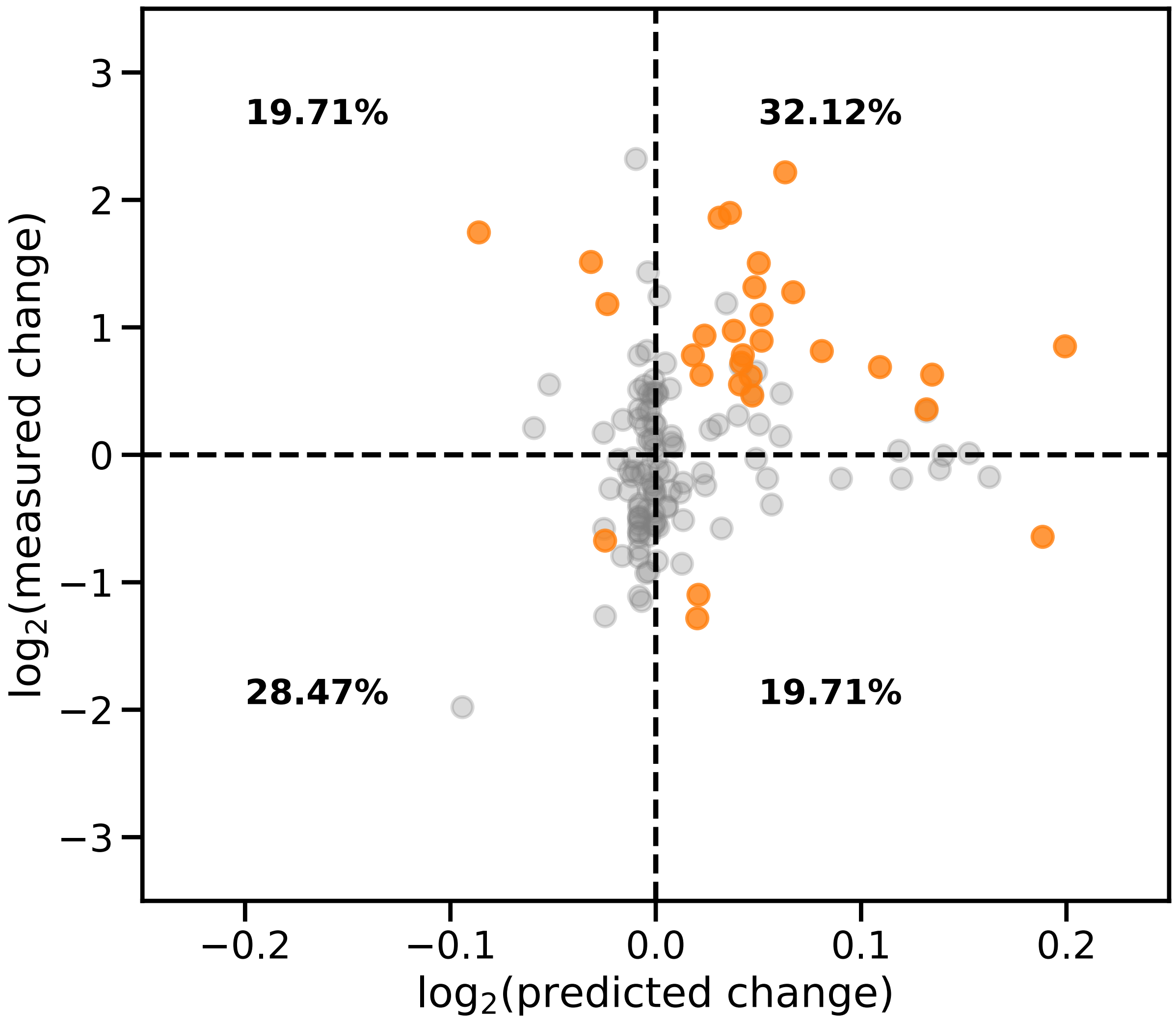


Figure S3: Comparison of predicted and measured changes in the abundance of 137 metabolites in UC patients relative to healthy controls. The dashed black lines separate the quadrants between correctly (upper right and lower left quadrants) and incorrectly predicted metabolites. Metabolites measured not to be significantly changed ($p > 0.05$) or predicted changes $dPR < \|0.014\|$ are shown in grey. All other metabolites are shown in orange. The percentage of all metabolites in each quadrant is shown.

### TABLES

Table S1: Predicted and measured metabolite changes corresponding to Figure 2b and S1a. Predictions were derived using RNA-Seq data from SLC25A51ko vs parental HEK293 cells). The columns list: names of measured metabolites, measured fold change, predicted fold change, respective standard deviations (5 replicates metabolomics, 3 replicates RNA-Seq).

| METABOLITE NAME | MEASURED | P-VALUE | MEASURED STD_DEV | PREDICTED | PREDICTED STD_DEV |
|---|---|---|---|---|---|
| **2,3-DIPHOSPHOGLYCERIC ACID** | 1.311 | 0.154 | 0.393 | 1.039 | 0.013 |
| **6-PHOSPHOGLUCONIC ACID** | 0.826 | 0.160 | 0.352 | 0.965 | 0.020 |
| **ADENOSINE DIPHOSPHATE** | 1.083 | 0.518 | 0.161 | 0.988 | 0.009 |
| **ADENOSINE MONOPHOSPHATE** | 1.408 | 0.011 | 0.310 | 0.982 | 0.008 |
| **ADENOSINE TRIPHOSPHATE** | 0.700 | 0.001 | 0.090 | 0.991 | 0.005 |
| **CITRIC ACID** | 0.644 | 0.001 | 0.100 | 0.815 | 0.046 |
| **CYCLIC AMP** | 0.838 | 0.053 | 0.140 | 1.018 | 0.030 |
| **D-GLUCOSE** | 0.648 | 0.000 | 0.075 | 1.029 | 0.084 |
| **D-SEDOHEPTULOSE 7-PHOSPHATE** | 0.666 | 0.001 | 0.117 | 0.953 | 0.163 |
| **FRUCTOSE 1,6-BISPHOSPHATE** | 1.068 | 0.605 | 0.223 | 0.871 | 0.036 |
| **FRUCTOSE 6-PHOSPHATE** | 1.488 | 0.016 | 0.389 | 1.029 | 0.080 |
| **FUMARIC ACID** | 4.964 | 0.000 | 0.703 | 0.756 | 0.070 |
| **GLUCOSE 1-PHOSPHATE** | 1.176 | 0.151 | 0.233 | 1.041 | 0.069 |
| **GLUCOSE 6-PHOSPHATE** | 0.839 | 0.157 | 0.378 | 1.091 | 0.027 |
| **GLYCINE** | 0.759 | 0.030 | 0.174 | 1.003 | 0.008 |
| **ISOCITRIC ACID** | 0.702 | 0.008 | 0.124 | 0.805 | 0.039 |
| **L-ALANINE** | 1.000 | 0.814 | 0.136 | 0.978 | 0.034 |
| **L-ARGININE** | 1.019 | 0.969 | 0.202 | 1.013 | 0.023 |
| **L-ASPARTIC ACID** | 1.862 | 0.000 | 0.310 | 1.016 | 0.018 |
| **L-GLUTAMIC ACID** | 0.482 | 0.000 | 0.057 | 1.055 | 0.018 |
| **L-GLUTAMINE** | 0.981 | 0.633 | 0.187 | 0.989 | 0.014 |
| **L-HISTIDINE** | 0.866 | 0.138 | 0.150 | 0.995 | 0.024 |
| **L-ISOLEUCINE** | 0.842 | 0.224 | 0.242 | 0.993 | 0.019 |
| **L-LACTIC ACID** | 1.228 | 0.049 | 0.207 | 1.050 | 0.028 |
| **L-LEUCINE** | 0.889 | 0.322 | 0.251 | 0.996 | 0.017 |
| **L-LYSINE** | 1.167 | 0.415 | 0.253 | 1.005 | 0.012 |
| **L-MALIC ACID** | 4.086 | 0.000 | 0.674 | 0.748 | 0.064 |
| **L-METHIONINE** | 0.664 | 0.075 | 0.245 | 1.008 | 0.050 |
| **L-PHENYLALANINE** | 1.045 | 0.965 | 0.228 | 1.030 | 0.047 |
| **L-PROLINE** | 0.396 | 0.000 | 0.069 | 0.944 | 0.030 |
| **L-SERINE** | 1.324 | 0.073 | 0.265 | 0.972 | 0.016 |
| **L-THREONINE** | 1.004 | 0.911 | 0.108 | 1.000 | 0.013 |
| **L-TRYPTOPHAN** | 1.156 | 0.902 | 0.462 | 1.003 | 0.031 |

| | | | | | |
|---|---|---|---|---|---|
| **L-TYROSINE** | 1.127 | 0.829 | 0.353 | 1.064 | 0.048 |
| **L-VALINE** | 0.906 | 0.346 | 0.241 | 0.994 | 0.020 |
| **OXOGLUTARIC ACID** | 1.304 | 0.045 | 0.260 | 0.963 | 0.026 |
| **PHOSPHOENOLPYRUVIC ACID** | 1.482 | 0.099 | 0.539 | 0.874 | 0.060 |
| **PYRUVIC ACID** | 2.401 | 0.000 | 0.999 | 1.026 | 0.015 |
| **SUCCINIC ACID** | 0.947 | 0.380 | 0.160 | 0.834 | 0.049 |
| **CIS-ACONITIC ACID** | 0.639 | 0.000 | 0.069 | 0.768 | 0.063 |
| **HYDROXYKYNURENINE** | 0.680 | 0.092 | 0.403 | 0.952 | 0.018 |
| **L-KYNURENINE** | 0.648 | 0.016 | 0.188 | 0.949 | 0.013 |
| **N-ACETYL-L-ASPARTIC ACID** | 0.558 | 0.001 | 0.139 | 0.974 | 0.011 |
| **NICOTINIC ACID** | 1.328 | 0.462 | 0.548 | 0.922 | 0.016 |
| **S-ADENOSYLMETHIONINE** | 0.482 | 0.010 | 0.204 | 1.002 | 0.003 |

Table S2: Predicted and measured metabolite changes using between UC patient and healthy controls. Predictions were based on the integration of differential proteomics data. The columns list: name of measured metabolites, measured fold change, level of significance (p-value) in measured changes, predicted fold change.

| METABOLITE NAME | MEASURED | P-VALUE | PREDICTED |
|---|---|---|---|
| **2-HYDROXY-ISOCAPROATE** | 0.730 | 0.459 | 0.999 |
| **3-HYDROXYHEXADECANOYLCARNITINE** | 2.488 | 0.001 | 1.034 |
| **3-HYDROXY-ISOVALERYL CARNITINE** | 1.719 | 0.005 | 1.030 |
| **3-HYDROXY-11Z-OCTADECENOYLCARNITINE** | 2.836 | 0.001 | 1.035 |
| **3-HYDROXY BUTYRYL CARNITINE** | 1.402 | 0.835 | 1.033 |
| **3-HYDROXYDODECANOYLCARNITINE** | 4.647 | 0.000 | 1.045 |
| **3-HYDROXY-TETRADECANOYL CARNITINE** | 2.421 | 0.001 | 1.047 |
| **GAMMA-AMINOBUTYRIC ACID** | 1.432 | 0.203 | 1.005 |
| **5-HYDROXYINDOLEACETIC ACID** | 2.700 | 0.000 | 0.997 |
| **5'-METHYLTHIOADENOSINE** | 0.763 | 0.372 | 1.040 |
| **PYROGLUTAMIC ACID** | 1.760 | 0.001 | 0.997 |
| **5-HYDROXYHEXANOIC ACID** | 0.701 | 0.506 | 1.009 |
| **7-HYDROXY-OCTANOATE** | 1.035 | 0.776 | 1.000 |
| **L-ACETYLCARNITINE** | 1.380 | 0.014 | 1.033 |
| **L-ARABINOSE** | 0.843 | 0.531 | 0.999 |
| **ARACHIDIC ACID** | 0.751 | 0.231 | 1.004 |
| **ARACHIDONIC ACID** | 1.086 | 0.296 | 0.997 |
| **L-ARGININE** | 0.879 | 0.506 | 1.038 |
| **L-ASPARAGINE** | 1.804 | 0.000 | 1.148 |
| **PELARGONIC ACID** | 0.254 | 0.635 | 0.937 |
| **PICOLINIC ACID** | 1.176 | 0.690 | 1.000 |
| **DECANOYLCARNITINE** | 3.632 | 0.004 | 1.022 |
| **L-OCTANOYLCARNITINE** | 3.726 | 0.002 | 1.025 |

| | | | |
|---|---|---|---|
| **(3S)-3-HYDROXYLINOLEOYL-COA** | 3.352 | 0.000 | 0.942 |
| **CYSTEINYLGLYCINE** | 1.068 | 0.690 | 1.005 |
| **DOCOSAPENTAENOIC ACID** | 1.165 | 0.279 | 0.996 |
| **DOCOSAPENTAENOIC ACID** | 1.459 | 0.077 | 0.996 |
| **L-CARNITINE** | 0.846 | 0.776 | 1.017 |
| **DOCOSAHEXAENOIC ACID** | 0.823 | 0.060 | 0.991 |
| **DODECANOIC ACID** | 0.914 | 0.506 | 0.993 |
| **LAUROYL CARNITINE** | 2.277 | 0.065 | 1.024 |
| **D-FRUCTOSE** | 2.270 | 0.004 | 0.984 |
| **FUMARIC ACID** | 1.210 | 0.190 | 0.989 |
| **D-GALACTOSE** | 0.577 | 0.098 | 0.989 |
| **L-GLUTAMINE** | 1.008 | 0.924 | 1.112 |
| **L-GLUTAMIC ACID** | 1.912 | 0.000 | 1.017 |
| **GUANOSINE** | 0.683 | 0.154 | 0.999 |
| **CIS-ACONITIC ACID** | 0.553 | 0.333 | 1.009 |
| **PALMITIC ACID** | 0.814 | 0.133 | 1.008 |
| **PALMITOLEIC ACID** | 1.045 | 0.582 | 1.006 |
| **HYPOXANTHINE** | 1.644 | 0.000 | 1.003 |
| **MYO-INOSITOL** | 0.822 | 0.065 | 1.005 |
| **INOSINE** | 1.107 | 0.459 | 1.043 |
| **ISOVALERYL CARNITINE** | 1.860 | 0.035 | 1.036 |
| **ISOVALERYL CARNITINE** | 2.142 | 0.013 | 1.036 |
| **LEUCYL-GLYCINE** | 0.530 | 0.011 | 0.998 |
| **LEUCYL-GLYCINE** | 1.396 | 0.014 | 0.998 |
| **L-LEUCINE** | 0.878 | 0.414 | 1.065 |
| **LINOLEIC ACID** | 0.525 | 0.000 | 0.997 |
| **LINOLEYL CARNITINE** | 1.157 | 0.144 | 0.960 |
| **ALPHA-LINOLENIC ACID** | 0.696 | 0.006 | 1.000 |
| **ALPHA-LINOLENYL CARNITINE** | 1.545 | 0.038 | 1.016 |
| **L-LYSINE** | 0.906 | 0.635 | 1.016 |
| **L-KYNURENINE** | 2.854 | 0.000 | 0.978 |
| **(Z)-13-OCTADECENOIC ACID** | 0.982 | 0.835 | 0.992 |
| **5,8,11-EICOSATRIENOIC ACID** | 0.916 | 0.482 | 0.991 |
| **TETRADECANOYLCARNITINE** | 1.620 | 0.091 | 1.029 |
| **L-MALIC ACID** | 1.178 | 0.333 | 1.021 |
| **N-ACETYL-L-ASPARTIC ACID** | 1.612 | 0.013 | 1.079 |
| **NIACINAMIDE** | 1.758 | 0.004 | 1.058 |
| **OLEIC ACID** | 0.627 | 0.000 | 0.983 |
| **OLEIC ACID** | 0.416 | 0.372 | 0.983 |
| **OLEOYLCARNITINE** | 1.272 | 0.177 | 0.997 |
| **ORNITHINE** | 1.718 | 0.012 | 1.013 |
| **LYSOPC(17:0)** | 0.463 | 0.000 | 0.994 |
| **LYSOPC(18:1(11Z))** | 0.597 | 0.026 | 0.994 |

| | | | |
|---|---|---|---|
| **LYSOPC(18:0)** | 0.702 | 0.012 | 0.994 |
| **PROPIONYLCARNITINE** | 1.396 | 0.077 | 1.043 |
| **L-PHENYLALANINE** | 0.925 | 0.894 | 1.101 |
| **L-PALMITOYLCARNITINE** | 1.127 | 0.531 | 0.983 |
| **PENTADECANOIC ACID** | 0.856 | 0.718 | 0.999 |
| **QUINOLINIC ACID** | 4.996 | 0.010 | 0.993 |
| **SQUALENE** | 0.919 | 0.247 | 1.001 |
| **L-SORBOSE** | 2.366 | 0.001 | 1.001 |
| **SUCCINIC ACID** | 0.668 | 0.065 | 0.983 |
| **SUCROSE** | 1.192 | 0.262 | 0.999 |
| **L-THREONINE** | 1.281 | 0.046 | 1.096 |
| **L-THREONINE** | 1.267 | 0.065 | 1.096 |
| **EICOSAPENTAENOIC ACID** | 0.451 | 0.001 | 0.995 |
| **L-TRYPTOPHAN** | 0.640 | 0.005 | 1.140 |
| **L-TYROSINE** | 0.996 | 0.864 | 1.102 |
| **URACIL** | 0.889 | 0.414 | 0.992 |
| **N-ACETYL-D-GLUCOSAMINE** | 0.831 | 0.065 | 0.985 |
| **S-ADENOSYLHOMOCYSTEINE** | 1.407 | 0.022 | 1.000 |
| **BETA-ALANINE** | 0.756 | 0.166 | 1.004 |
| **BUTYRYLCARNITINE** | 1.532 | 0.021 | 1.033 |
| **TIGLYL CARNITINE** | 1.651 | 0.009 | 1.029 |
| **GLUTARYL CARNITINE** | 0.411 | 0.022 | 1.014 |
| **HEXANOYL CARNITINE** | 1.963 | 0.026 | 1.027 |
| **2-OCTENOYLCARNITINE** | 0.979 | 0.835 | 1.034 |
| **CITRIC ACID** | 1.179 | 0.144 | 1.036 |
| **CITRULLINE** | 1.466 | 0.026 | 1.029 |
| **CREATINE** | 0.765 | 0.098 | 0.994 |
| **CREATININE** | 0.921 | 0.985 | 0.997 |
| **GLUCOSE-1,3-MANNOSE OLIGOSACCHARIDE** | 0.670 | 0.414 | 1.022 |
| **D-GLUCOSE** | 0.644 | 0.115 | 0.998 |
| **GLUTAMYL-LEUCINE** | 1.275 | 0.046 | 0.998 |
| **GLYCEROL 3-PHOSPHATE** | 0.467 | 0.003 | 1.015 |
| **PALMITOLEOYL-CARNITINE** | 1.464 | 0.175 | 0.965 |
| **HEPTADECANOIC ACID** | 1.083 | 0.805 | 0.998 |
| **ISOCITRIC ACID** | 1.238 | 0.091 | 1.028 |
| **L-ISOLEUCINE** | 0.878 | 0.436 | 1.086 |
| **ISOLECYL-ASPARTATE** | 1.358 | 0.091 | 0.999 |
| **D-MALTOSE** | 0.675 | 0.352 | 1.001 |
| **L-METHIONINE** | 0.886 | 0.776 | 1.119 |
| **STEARIC ACID** | 1.107 | 0.032 | 1.005 |
| **OCTADECADIENOATE (N-C18:2)** | 0.802 | 0.217 | 1.000 |
| **1-EICOSADIENOYLGLYCEROPHOSPHOCHOLINE (DELTA 11,14)** | 1.281 | 0.333 | 0.994 |
| **1-LINOLEOYLGLYCEROPHOSPHOCHOLINE (DELTA 9,12)** | 0.573 | 0.000 | 0.994 |

| | | | |
|---|---|---|---|
| **1-PENTADECANOYLGLYCEROPHOSPHOCHOLINE, SN1-LPC (15:0)** | 0.683 | 0.032 | 0.994 |
| **1-OCTADECA-TRIENOYLGLYCEROPHOSPHOCHOLINE, SN1-LPC (18:3, DELTA 9, 12, 15)** | 0.657 | 0.024 | 0.994 |
| **1-EICOSENOYLGLYCEROPHOSPHOCHOLINE (DELTA 11) ,SN1-LPC (20:1)** | 0.714 | 0.106 | 0.994 |
| **1-DIHOMO-LINOLENOYLGLYCEROPHOSPHOCHOLINE (20:3, DELTA 8, 11, 14), LYSOPC A C20:3** | 1.425 | 0.008 | 0.994 |
| **LYSOPC(20:4(8Z,11Z,14Z,17Z))** | 1.719 | 0.000 | 0.994 |
| **1-EICOSAPENTENOYLGLYCEROPHOSPHOCHOLINE (DELTA 5, 8, 11, 14, 17), SN1-LPC (20:5)** | 0.640 | 0.077 | 0.994 |
| **1-DOCOSAPENTENOYLGLYCEROPHOSPHOCHOLINE (DELTA 7, 10, 13, 16, 19), SN1-LPC (22:5)-W3** | 1.217 | 0.091 | 0.994 |
| **1-DOCOSAHEXENOYLGLYCEROPHOSPHOCHOLINE (DELTA 4, 7, 10, 13, 16, 19), SN1-LPC (22:6)** | 0.711 | 0.019 | 0.994 |
| **1-PALMITOYLGLYCEROPHOSPHOCHOLINE** | 0.656 | 0.000 | 0.994 |
| **1-PALMITOLEOYLGLYCEROPHOSPHOCHOLINE (DELTA 9)** | 0.749 | 0.203 | 0.994 |
| **1-HEPTADECANOYLGLYCEROPHOSPHOETHANOLAMINE (C17:0 PE)** | 0.821 | 0.314 | 1.000 |
| **1-EICOSATRIENOYLGLYCEROPHOSPHOETHANOLAMINE (DELTA 11, 14, 17), LPE (20:3)** | 0.681 | 0.060 | 1.000 |
| **1-ARACHIDONOYL-SN-GLYCERO-3-PHOSPHOETHANOLAMINE** | 1.389 | 0.024 | 1.001 |
| **PECTIN** | 1.095 | 0.414 | 0.999 |
| **1-PALMITOYLGLYCEROPHOSPHOETHANOLAMINE** | 0.561 | 0.001 | 1.001 |
| **ALPHA-N-PHENYLACETYL-L-GLUTAMINE** | 1.404 | 0.879 | 0.999 |
| **PHENYLALANYL-LEUCINE** | 0.743 | 0.955 | 0.997 |
| **PHENYLALANYL-PHENYLALANINE** | 0.824 | 0.648 | 0.998 |
| **L-PROLINE** | 1.022 | 0.231 | 1.086 |
| **D-RIBOSE** | 1.505 | 0.014 | 0.999 |
| **O-OCTADECANOYL-R-CARNITINE** | 1.147 | 0.333 | 1.019 |
| **SUBERIC ACID** | 0.859 | 0.608 | 1.009 |
| **TETRADECENOYL CARNITINE** | 1.572 | 0.144 | 1.034 |
| **MYRISTIC ACID** | 0.902 | 0.333 | 0.995 |
| **L-VALINE** | 1.547 | 0.024 | 1.098 |
| **16-HYDROXY HEXADECANOIC ACID** | 0.914 | 0.547 | 1.004 |
| **OMEGA HYDROXY TETRADECANOATE (N-C14:0)** | 0.979 | 0.835 | 1.000 |
| **D-XYLOSE** | 0.973 | 0.690 | 0.988 |
